# Toward Standardized Ex Vivo Joint Models: Impact of Glucose and Oxygen Levels for Enhanced Tissue Maintenance

**DOI:** 10.64898/2026.02.14.704322

**Authors:** Fatemeh Safari, Jovana Zvicer, Sibylle Grad, Martin J. Stoddart, Zhen Li

## Abstract

Ex vivo models bridge *in vitro* and *in vivo* systems by preserving native extracellular matrix architecture and multicellular interactions. In articular joint research, osteochondral–synovial co-cultures are particularly valuable for studying bone–cartilage crosstalk and synovial inflammatory regulation. However, a lack of standardized culture conditions regarding glucose and oxygen, two key regulators of cellular metabolism, limits reproducibility and translational relevance. This study aims to define how glucose and oxygen conditions influence joint tissues maintenance in an *ex vivo* model. Bovine osteochondral explants and synovium are harvested from the stifle joint and co-cultured using either high glucose DMEM (HG, 4.5 g/L) or low glucose DMEM (LG, 1 g/L) under hyperoxic (21% O_2_) or physioxic (5% O_2_) conditions. Cell viability, gene expression, and metabolomic profiles are evaluated across tissues. LG conditions increase cell death in the deep zone of cartilage and in subchondral bone. Gene expression and metabolomic analyses reveal tissue-specific effects of glucose and oxygen. In cartilage and bone, glucose-dependent gene regulation and metabolic changes occur under hyperoxia but are largely absent under physioxia, indicating buffering of glucose responses. Gene-specific sensitivity to glucose and oxygen is observed in bone and synovium; however, glucose-induced metabolic responses persist under physioxia only in synovium. Overall, these findings identify oxygen and glucose as critical modulators of joint tissue physiology and support the use of HG, physioxic culture conditions to improve cell viability and stabilize molecular outcomes in *ex vivo* joint models. This optimized ex vivo model provides platforms for investigating mechanisms relevant to joint-related diseases.

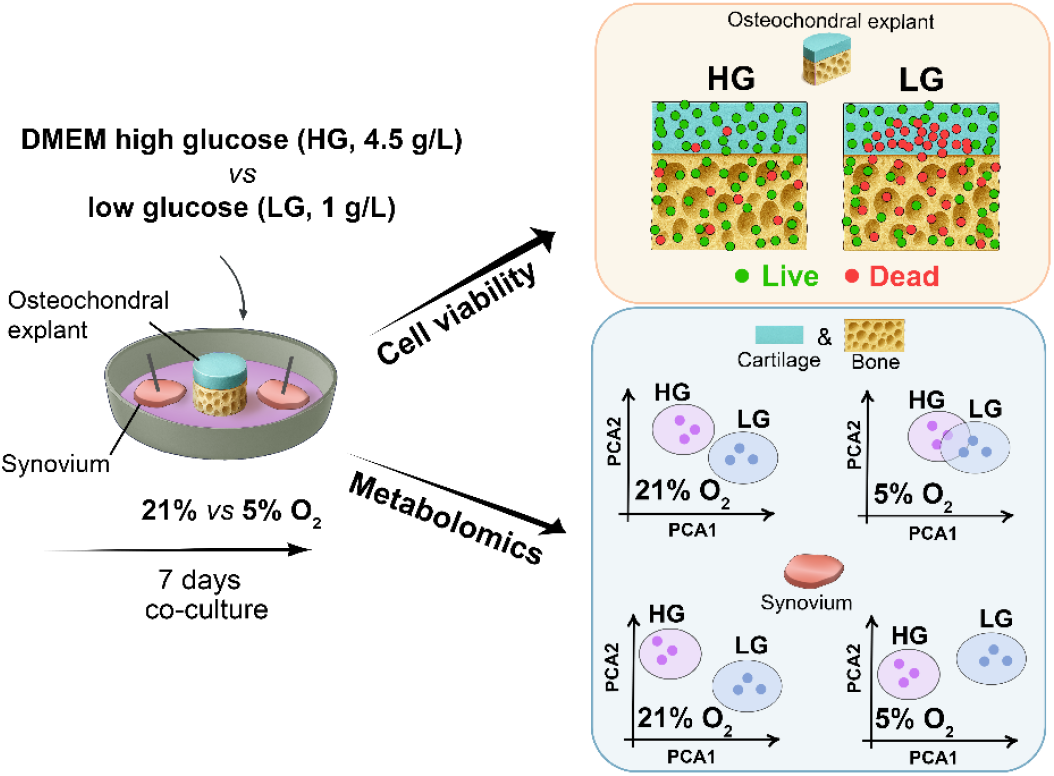

This study evaluates the effects of glucose concentration and oxygen tension in an ex vivo joint co-culture system to define optimal culture conditions. High glucose medium and physioxic conditions support tissue viability, preserve homeostasis, and enhance the physiological relevance of the ex vivo model.

## 1 INTRODUCTION

*Ex vivo* explant-based models occupy a unique position between simplified *in vitro* systems and complex *in vivo* animal studies [1]. By preserving native extracellular matrix (ECM) architecture and maintaining authentic cell–cell and cell–matrix interactions, they capture biological features that are lost in monolayer cultures or even in advanced 3D constructs such as pellets or organoid models [2, 3].

For articular cartilage regeneration, *ex vivo* models have been widely applied in the functional assessment of emerging therapies [4-7]. Beyond regenerative applications, these systems play a central role in osteoarthritis (OA) research, allowing investigations of cartilage degeneration, enabling controlled disease-like environments, and helping in pre-screening of therapeutic candidates [8].

To create an *ex vivo* environment that more accurately reflects *in vivo* joint biology and enhances the translational value of preclinical findings, the most reliable strategy is to use osteochondral explants in combination with synovial tissue [1]. Osteochondral explants already outperform cartilage only models, as the presence of subchondral bone markedly alters inflammatory and catabolic signaling and improves chondrocyte survival, likely through the release of bone-derived mediators [9]. Including synovial tissue further enhances the physiological relevance of the model, as recent studies have demonstrated that synovial tissue macrophages (STMs) form an immune-protective lining barrier that regulates joint inflammation and may control disease progression or remission in inflammatory joint disorders, such as rheumatoid arthritis [10].

Although this integrated osteochondral–synovium model provides a robust foundation for physiologically relevant *ex vivo* joint models, the field still lacks standardized protocols for establishing such models. This gap is particularly evident in the definition of two fundamental culture parameters, glucose availability and oxygen tension, which are critical for maintaining tissue physiology. It is generally recognized that all three joint tissues operate under markedly reduced oxygen conditions, with cartilage experiencing gradients from approximately 6% in the superficial zone to as low as 1% in the deep zone [11]. Vascularized tissues such as bone and synovium exhibit higher levels of oxygen between 5% to 12.5% [12]. Physiological glucose concentrations in cartilage, bone and synovial fluid are substantially lower than those present in standard high glucose (4.5 g/L, ∼25 mM) media and are typically around 5 mM (∼0.9 g/L) [13, 14], which is in the range of normal blood glucose.

However, studies have shown contrasting results regarding tissue metabolism and its maintenance *in vitro*, depending on whether physiological conditions (∼5 mM glucose and 5% O_2_ physioxia) or non-physiological conditions (∼25 mM glucose and 21% O_2_ hyperoxia) were used. It is shown that physioxia promotes cartilage matrix synthesis by enabling direct binding of HIF-2α to a regulatory site in SOX9 gene, while suppressing cartilage degradation through predominantly HIF-1α–mediated anti-catabolic responses [15]. In bone tissue, low oxygen levels exert a dual HIF-dependent effect, where HIF-1α promotes anabolic responses while on contrary HIF-2α limits further catabolic effects by inhibiting osteoblast differentiation and bone mineralization [16]. These findings are further supported by an *ex vivo* study using porcine osteochondral explants, which demonstrated enhanced cartilaginous matrix deposition in the defect site under 2% oxygen [17]. Furthermore, an *in vivo* study in rats maintained under 12% oxygen conditions demonstrated enhanced cartilage matrix formation and improved structural organization within cartilage defects after 4 weeks, compared with animals maintained under 21% oxygen[18].

Regarding glucose availability, concentrations above the physiological level (5 mM) had adverse effects on synovium, cartilage, and bone. In synovium explants, high glucose (12.5 mM) reduced DNA content, collagen levels, and GAG deposition compared to physiological concentrations (5 mM) [19]. In addition, in bone, acute exposure to high glucose (25 mM) enhanced glycolysis and oxidative phosphorylation, and chronic exposure resulted in uncoupled oxygen consumption, increased ROS generation, downregulation of key mitochondrial bioenergetic proteins, and consequently elevated oxidative stress [20]. In cartilage, high glucose reduced collagen and GAG content as well as cilia length of chondrocytes [19]. Despite these adverse effects, different studies have demonstrated that standard laboratory culture conditions, typically employing high glucose DMEM (25 mM) and atmospheric oxygen levels (21% O_2_), can maintain cartilage viability and structural integrity even during long-term culture [21].

Given the lack of standardized protocols and the absence of data-driven guidance on optimal oxygen and glucose concentrations for *ex vivo* models of the articular joint, in this study, osteochondral explants co-cultured with synovial tissue were evaluated in two commonly used culture media, DMEM low glucose (LG, 1 g/L, ∼5.6 mM) and high glucose (HG, 4.5 g/L, ∼25 mM) and exposed to either 21% oxygen (hyperoxic) or 5% oxygen (physioxic) (Figure 1). By comparing cell viability, gene expression, and metabolomic profiles across cartilage, bone, and synovium, we aimed to identify culture conditions that maintain joint tissue viability and cell phenotype *ex vivo* and support physiologically relevant metabolic activity, thereby providing evidence-based recommendations for future joint research using *ex vivo* models.

**Figure 1:**
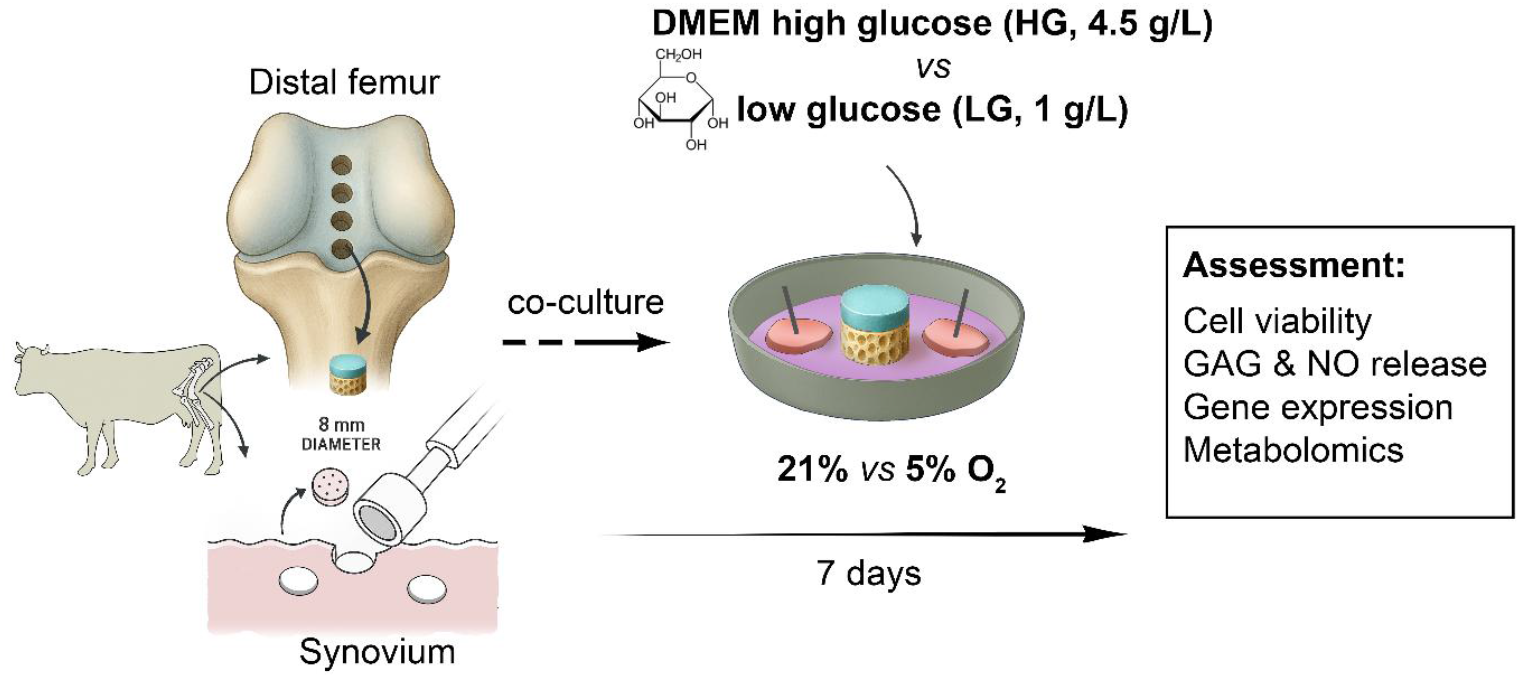
Schematic representation of experimental design.

## 2 RESULTS

### 2.1 Cell viability

Cell viability in cartilage, subchondral bone, and synovium tissues was assessed separately after 7 days of co-culture in either HG or LG medium and under two oxygen tensions (21% O_2_, considered hyperoxic; 5% O_2_, considered physioxic).

In cartilage, cell viability was assessed using LDH/ethidium homodimer staining. Viability varied across the cartilage depth and between the central and peripheral regions of the explants. In the peripheral region, cells were predominantly viable; therefore, quantitative evaluation of cell viability was focused on the central region of the explant, where the greatest variability was observed microscopically (Figure 2A). In the superficial zone (∼0–500 µm), no differences in the cell viability were observed between different glucose or oxygen conditions. In the middle zone (∼500–1000 µm), LG medium increased the number of dead cells compared to HG medium, although this difference was only significant under 5% O_2_ in the 800–1000 µm region (71 vs. 3 dead cells/area, *p* < 0.01, Figure 2A, B). In the deep zone (∼1000–1300 µm), LG similarly resulted in higher cell death in both oxygen conditions (94 and 128 dead cells/area, in LG-21% and LG-5% O_2_ vs. 6 and 8 dead cells/area, in HG-21% and HG-5% O_2_, respectively; *p* < 0.0001). In the calcified zone (∼1300–1500 µm), cell death tended to be higher under LG than HG in both oxygen levels, although there was no significant difference due to donor variability. Overall, the average number of dead cells/area, across cartilage depth was higher in LG than HG under both oxygen tensions (LG: 41 at 21% O_2_ and 57 at 5% O_2_; HG: 8 at 21% O_2_ and 10 at 5% O_2_; *p* < 0.05 for 21% O_2_ and *p* < 0.01 for 5% O_2_; Figure 2C).

**Figure 2.**
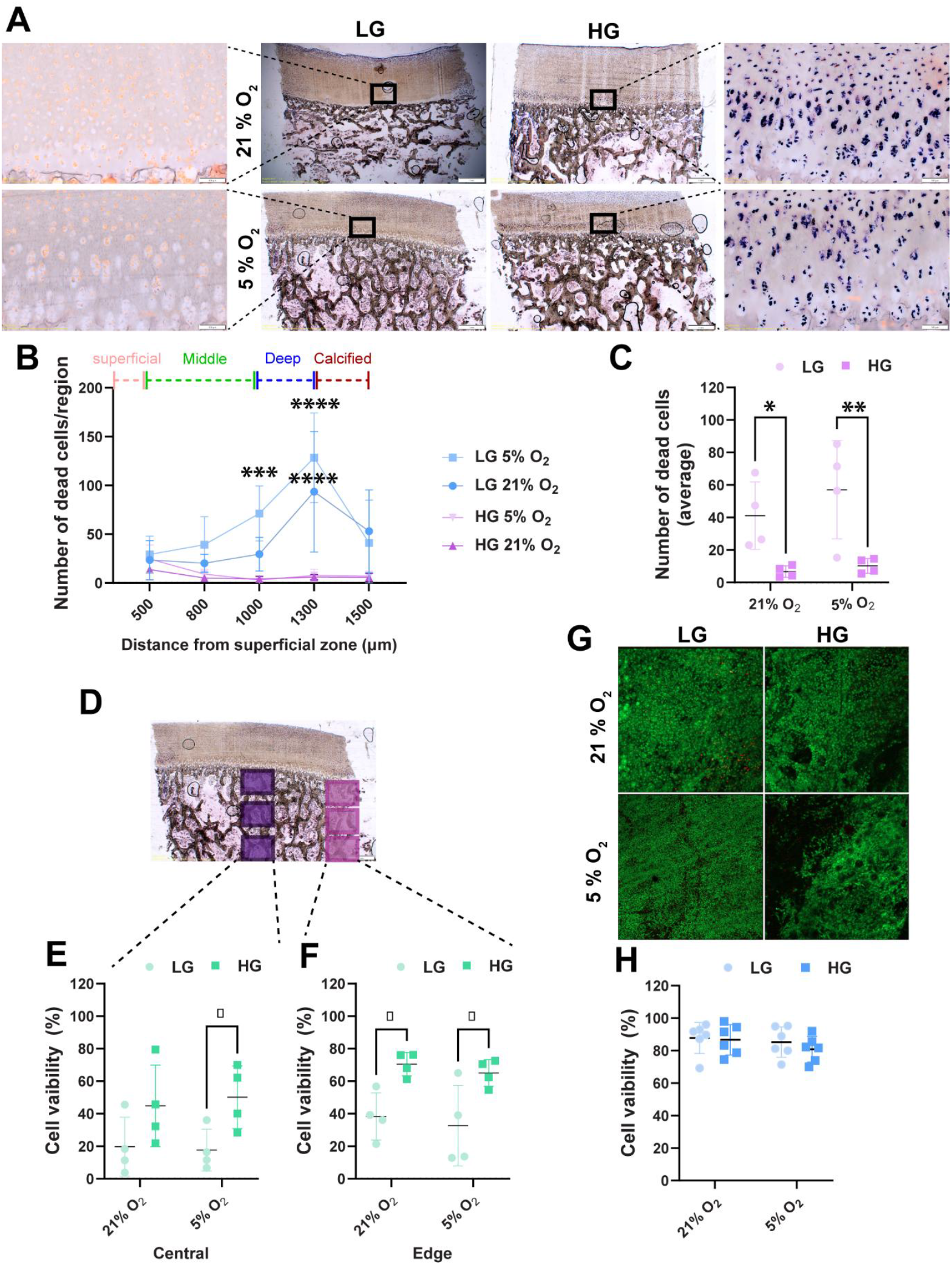
Cell viability of joint tissues. Co-cultures of osteochondral and synovium tissues were performed for 7 days in low glucose (LG) and high glucose (HG) DMEM under physioxia (5% O_2_) and hyperoxia (21% O_2_). (A) LDH/Ethidium homodimer staining in osteochondral tissues. (B) Average number of dead cells per counted area in different regions of cartilage (significancy is calculated between LG and HG either under 5% or 21% O_2_), and (C) in full thickness of cartilage. (D-F) Cell viability in subchondral bone was evaluated according to the pattern in figure (D) after LDH/Ethidium homodimer staining in (E) the central region and (F) the peripheral region of the subchondral bone. (G) Calcein/ethidium homodimer staining was performed for evaluation of cell viability in (H) synovium tissue. Results are presented as means of 4 donors for cartilage and bone and 6 donors for synovium. Statistical analysis was performed using 2-way ANOVA, with significance levels indicated as *p < 0.05, **p < 0.01, ****p < 0.0001.

In subchondral bone, LDH/ethidium homodimer staining also indicated greater cell viability at the explant periphery than in the central region. In the central region, cell viability was higher in HG than LG, reaching significance only under 5% O_2_ (50% vs. 18% viability, *p* < 0.05; Figure 2E). At the explant edge, LG significantly reduced cell viability compared with HG under both oxygen conditions (LG: 38% at 21% O_2_ and 33% at 5% O_2_; HG: 70% at 21% O_2_ and 65% at 5% O_2_; *p* < 0.05; Figure 2F).

In synovium tissue, calcein/ethidium homodimer staining showed that synovial cell viability was not affected by glucose or oxygen levels. Cell viability remained high across all conditions: 88% (LG-21% O_2_), 85% (LG-5% O_2_), 87% (HG-21% O_2_), and 81% (HG-5% O_2_; Figure 2G, H).

### 2.2 GAG and NO release from whole-joint tissues

GAG and NO levels in conditioned media were quantified over the 7-day culture period (Figure 3A and 3B, cumulative release). The release of GAG and NO was not significantly influenced by glucose concentration or oxygen tension. Consistently, Safranin O/Fast Green staining confirmed comparable proteoglycan staining intensity across culture conditions (Figure 3C).

**Figure 3.**
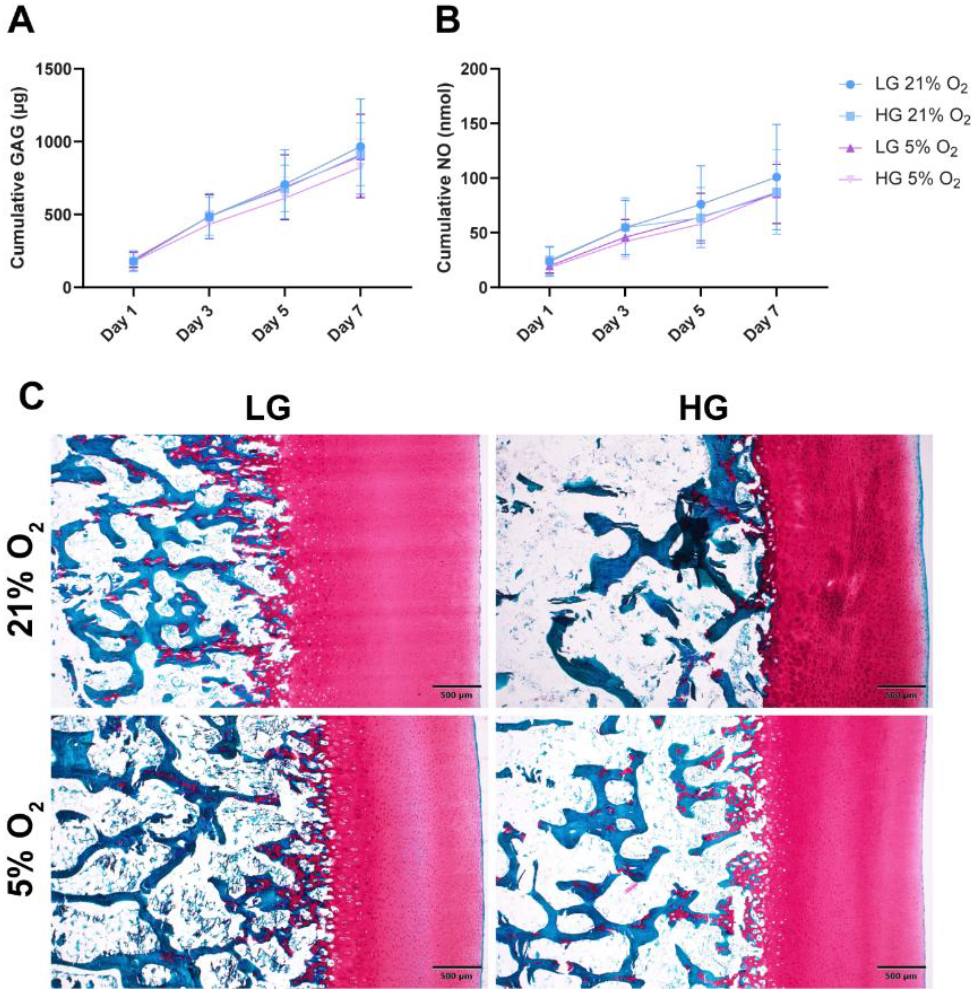
GAG and NO release. Cumulative release of (A) GAG and (B) NO during 7 days of culture in the presence of low glucose (LG) and high glucose (HG) DMEM under physioxia (5% O_2_) and hyperoxia (21% O_2_) conditions. (C) Safranin O/Fast Green staining of osteochondral tissues after 7 days of culture. Results are presented as means of 4 replicates from 5 donors for every tissue (n=20). Statistical analysis was performed using 2-way ANOVA.

### 2.3 Gene expression in individual joint tissues: cartilage, bone, and synovium

RT-qPCR analysis was performed separately on cartilage, subchondral bone, and synovium after 7 days of co-culture for different experimental conditions. Respective tissues on day 0 served as controls for all analyses.

In cartilage, *ACAN* transcript levels were upregulated in HG compared with LG under 21% O_2_ (p < 0.05), whereas no difference was detected under 5% O_2_ (Figure 4A). *COL2* and *COMP* expressions were reduced in LG under 21% O_2_ compared with LG under 5% O_2_; however, these transcripts were not affected by glucose concentration (Figure 4B and 4C, respectively). *COL10, ADAMTS4, MMP13*, and *IL6* transcripts were all upregulated in HG compared with LG under 21% O_2_ (p < 0.01, 0.001, 0.01, and 0.05, respectively) (Figure 4D, 4E, 4F and 4G, respectively). These differences were not observed under 5% O_2_, as transcript levels in HG were significantly lower at 5% O_2_ than at 21% O_2_ (p < 0.0001, 0.05, 0.0001, and 0.001 for *COL10A1, ADAMTS4, MMP13*, and *IL6*, respectively). *IL8* expression was not affected by either glucose or oxygen concentration (Figure 4H).

**Figure 4.**
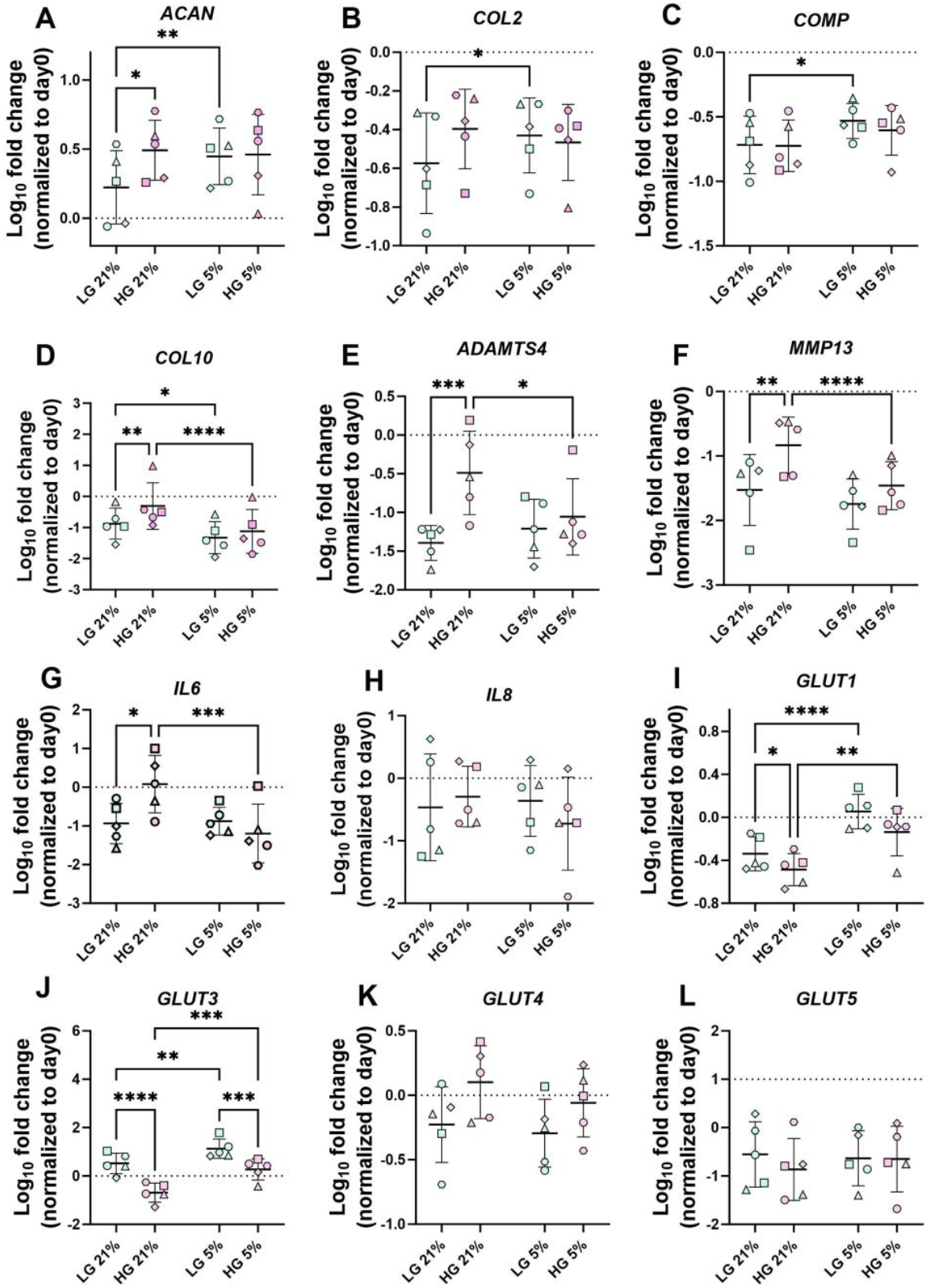
Gene expression levels in cartilage tissue cultured for 7 days in low glucose (LG) and high glucose (HG) DMEM under physioxia (5% O2) and hyperoxia (21% O_2_) conditions: (A) *ACAN*, (B) *COL2*, (C) *COMP*, (D) *COL10*, (E) *ADAMTS4*, (F) *MMP13*, (G) *IL6*, (H) *IL8*, (I) *GLUT1*, (J) *GLUT3*, (K) *GLUT4*, and (L) *GLUT5*. Data is normalized to fresh tissue collected on day 0. Results are presented as means of three replicates per donor (n = 5 donors, depicted with different symbols for each donor). Statistical analysis was performed using 2-way ANOVA, with significance levels indicated as *p < 0.05, **p < 0.01, ***p < 0.001, ****p < 0.0001.

Expression of glucose transporters *GLUT1, GLUT2, GLUT3, GLUT4*, and *GLUT5* was also evaluated. *GLUT2* transcripts were not detected in cartilage. *GLUT1* and *GLUT3* were upregulated in LG compared with HG under both oxygen conditions (Figure 4I and 4J respectively); for *GLUT1*, this difference did not reach significance at 5% O_2_. 5% O_2_ increased the expression of both transporters in LG and HG (*GLUT1*: p < 0.0001 and 0.001; *GLUT3*: p < 0.01 and 0.001 in LG and HG, respectively). *GLUT4* and *GLUT5* transcript levels were not affected by glucose or oxygen concentration (Figure 4K and 4L, respectively).

In subchondral bone, *ALPL* transcript levels were downregulated under 5% O_2_ compared with 21% O_2</sub>_ in both LG and HG conditions (p < 0.05 and p < 0.001, respectively; Figure 5A). *COL1* and *IL8* expressions were not affected by either glucose concentration or oxygen tension (Figure 5B and 5E, respectively). *BGLAP* and *IL6* expression was insensitive to glucose levels but was reduced under 21% O_2_ compared with 5% O_2_ (p < 0.01 and p < 0.05, respectively; Figure 5C and 5F, respectively). *VEGF* transcript levels were upregulated in LG compared with HG under 5% O_2_ (p < 0.01) and were also elevated under 5% O_2_ compared with 21% O_2_ within the LG condition (p < 0.05; Figure 5D). Analysis of glucose transporter gene expression revealed that *GLUT1* was downregulated in HG compared with LG at both 21% and 5% O_2_ (p < 0.01, Figure 5G). In addition, *GLUT1* expression increased in HG under 5% O_2_ relative to 21% O_2_ (p < 0.05). *GLUT3* was downregulated in HG compared with LG only under 5% O_2_ (p < 0.01, Figure 5H). *GLUT4* was upregulated in HG relative to LG under 21% O_2_, and this increase was diminished under 5% O_2_ (p < 0.001, Figure 5I). *GLUT5* expression was not influenced by glucose concentration or oxygen tension (Figure 5J).

**Figure 5.**
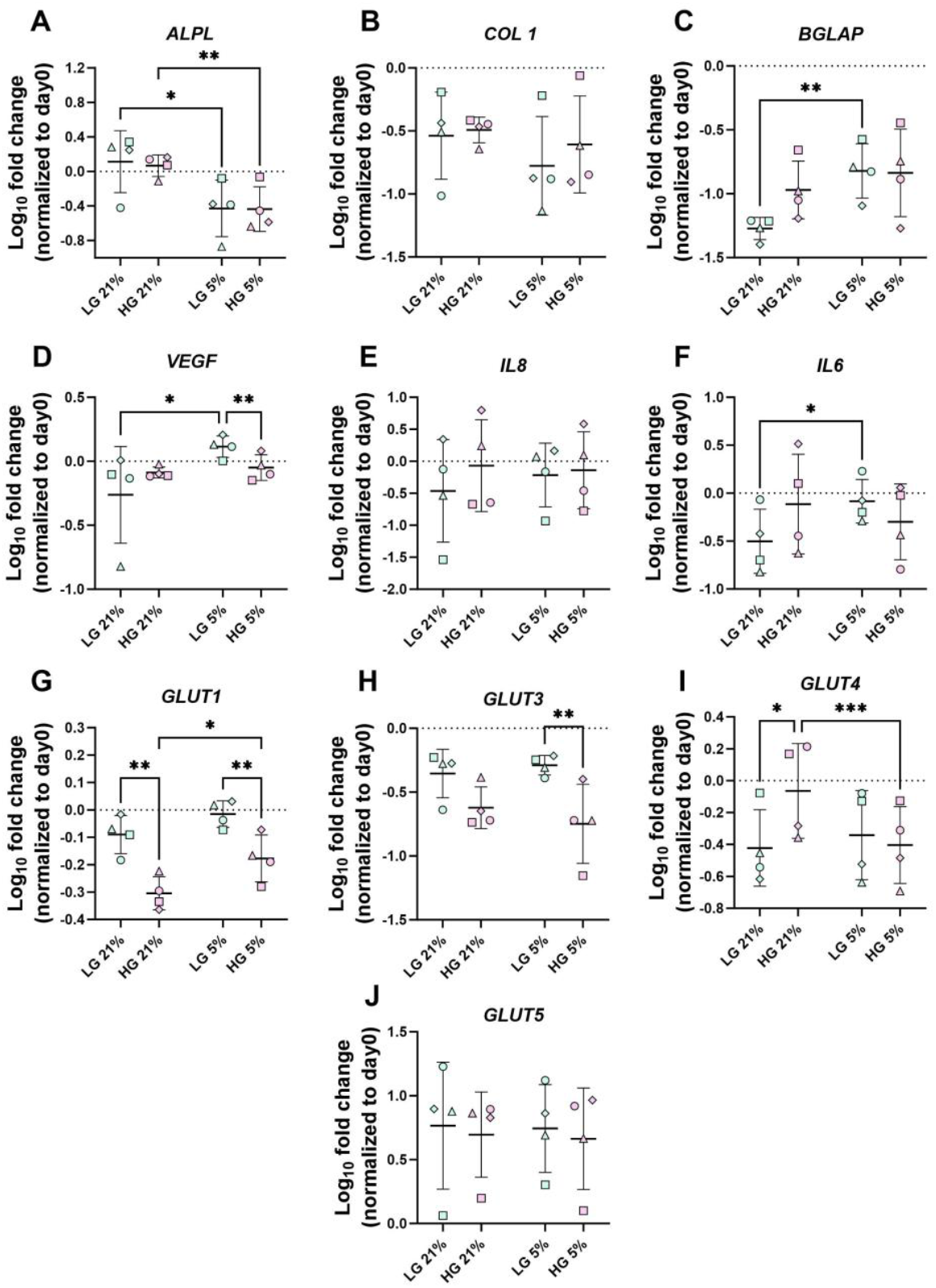
Gene expression levels in subchondral bone tissue cultured for 7 days in low glucose (LG) and high glucose (HG) DMEM under physioxia (5% O_2_) and hyperoxia (21% O_2_) conditions: (A) ALP, (B) COL1, (C) BGLAP, (D) VEGF, (E) IL8, (F) IL6, (G) GLUT1, (H) GLUT3, (I) GLUT4, and (J) GLUT5. Data is normalized to fresh tissue collected on day 0. Results are presented as means of three replicates per donor (n = 4 donors, depicted with different symbols for each donor). Statistical analysis was performed using 2-way ANOVA, with significance levels indicated as *p < 0.05, **p < 0.01, ***p < 0.001, ****p < 0.0001.

In synovium, *VEGF, MMP3*, and *IL8* transcript levels were not affected by glucose concentration or oxygen tension (Figure 6A, 6C, and 6F, respectively). *ADAMTS5* and *MMP1* showed differential sensitivity to oxygen: *ADAMTS5* was downregulated in LG condition compared to HG under both 21% and 5% O_2_ (p < 0.01 in 21% O_2_; p < 0.001 in 5% O_2_ Figure 6B), whereas *MMP1* was downregulated under 5% O_2_ in both glucose conditions (p < 0.05, Figure 6D, respectively). *IL6* expression was increased in HG compared with LG, but only under 21% O_2_ (p < 0.0001, Figure 6E). Analysis of glucose transporters showed that *GLUT1* expression was unaffected by glucose or oxygen (Figure 6G), and *GLUT2* was not detectable in synovium. *GLUT3* expression was reduced in HG compared with LG under 21% O_2_ (p < 0.05, Figure 6H), and this difference was diminished under 5% O2. *GLUT4* expression was higher in HG than LG under 21% O_2_ (p < 0.05, Figure 6I), with no differences observed at 5% O_2_. *GLUT5* expression was decreased under 5% O_2_ compared with 21% O_2_, but only in LG (p < 0.05, Figure 6J).

**Figure 6.**
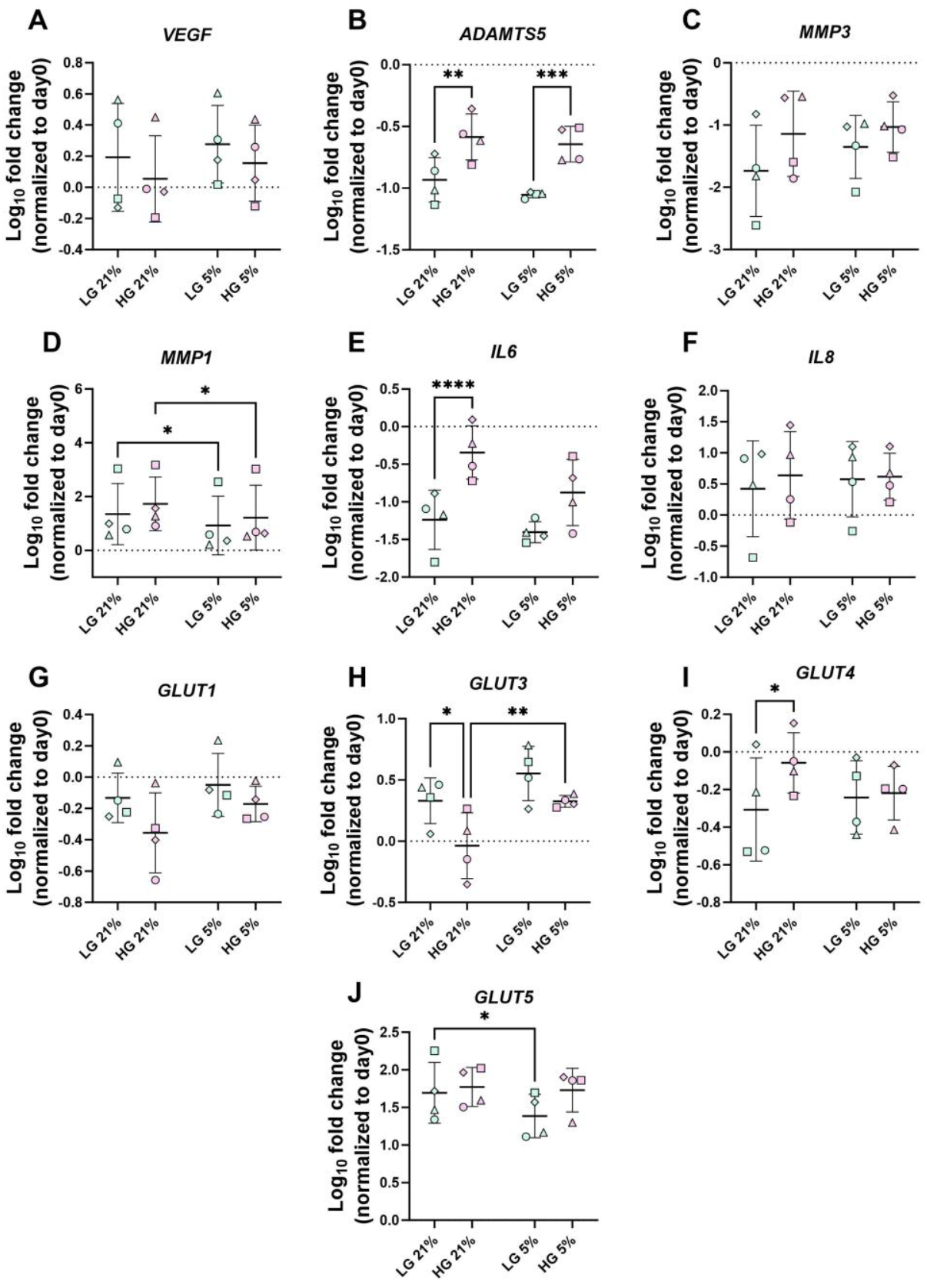
Gene expression levels in synovium tissue cultured for 7 days in low glucose (LG) and high glucose (HG) DMEM under physioxia (5% O_2_) and hyperoxia (21% O_2_) conditions: (A) *VEGF*, (B) *ADAMTS5*, (C) *MMP3*, (D) *MMP1*, (E) *IL6*, (F) *IL8*, (G) *GLUT1*, (H) *GLUT3*, (I) *GLUT4*, and (J) *GLUT5*. Data is normalized to fresh tissue collected on day 0. Results are presented as means of three replicates per donor (n = 4 donors, depicted with different symbols for each donor). Statistical analysis was performed using 2-way ANOVA, with significance levels indicated as *p < 0.05, **p < 0.01, ***p < 0.001, ****p < 0.0001.

### 2.4 Metabolomics in individual joint tissues: cartilage, bone, and synovium

To examine the metabolomic profiles of tissues cultured under varying glucose and oxygen levels, we performed untargeted metabolomics on cartilage, subchondral bone, and synovium separately. Fresh tissue from each compartment was processed in parallel and used as a control. In total, 121 metabolites were detected across cartilage, bone, and synovium.

Principal component analysis (PCA) revealed a clear separation between fresh tissues and cultured tissues for 7 days (Figure 7A, B, C), a pattern observed consistently across all three tissue types. To evaluate differences associated with the culture variables themselves, PCA was subsequently performed on all cultured samples without the control group (Supplementary material, Figure S1). To evaluate the effects of glucose and oxygen levels specifically, PCA was then repeated for each of the two conditions, separately. In cartilage, under 21% O_2_, HG and LG samples showed modest separation with partial overlap, whereas under 5% O_2_ no clustering between the two glucose conditions was observed. When comparing oxygen levels, a slight distinction between 21% and 5% O_2_ was evident in the HG groups (PC1: 39.1%), while LG samples showed no clear separation (Figure 7A). In subchondral bone, HG and LG conditions showed a clear separation under 21% O_2_, primarily along PC2 (20.5%). This clustering was not observed under 5% O_2_. Comparison of oxygen levels revealed a strong separation between 21% and 5% O_2_ within the LG groups along PC2 (23.3%), whereas HG samples exhibited only a weak distinction (Figure 7B). Synovial tissue showed the strongest glucose-driven effects. HG and LG samples formed distinct clusters under both oxygen levels, with separation occurring mainly along PC1 (42.8% at 21% O_2_; 38.5% at 5% O_2_). When comparing oxygen levels, HG samples showed only slight clustering between 21% and 5% O_2_ (PC1: 36%), while LG samples demonstrated clearer separation (PC1: 36.8%, Figure 7C).

**Figure 7.**
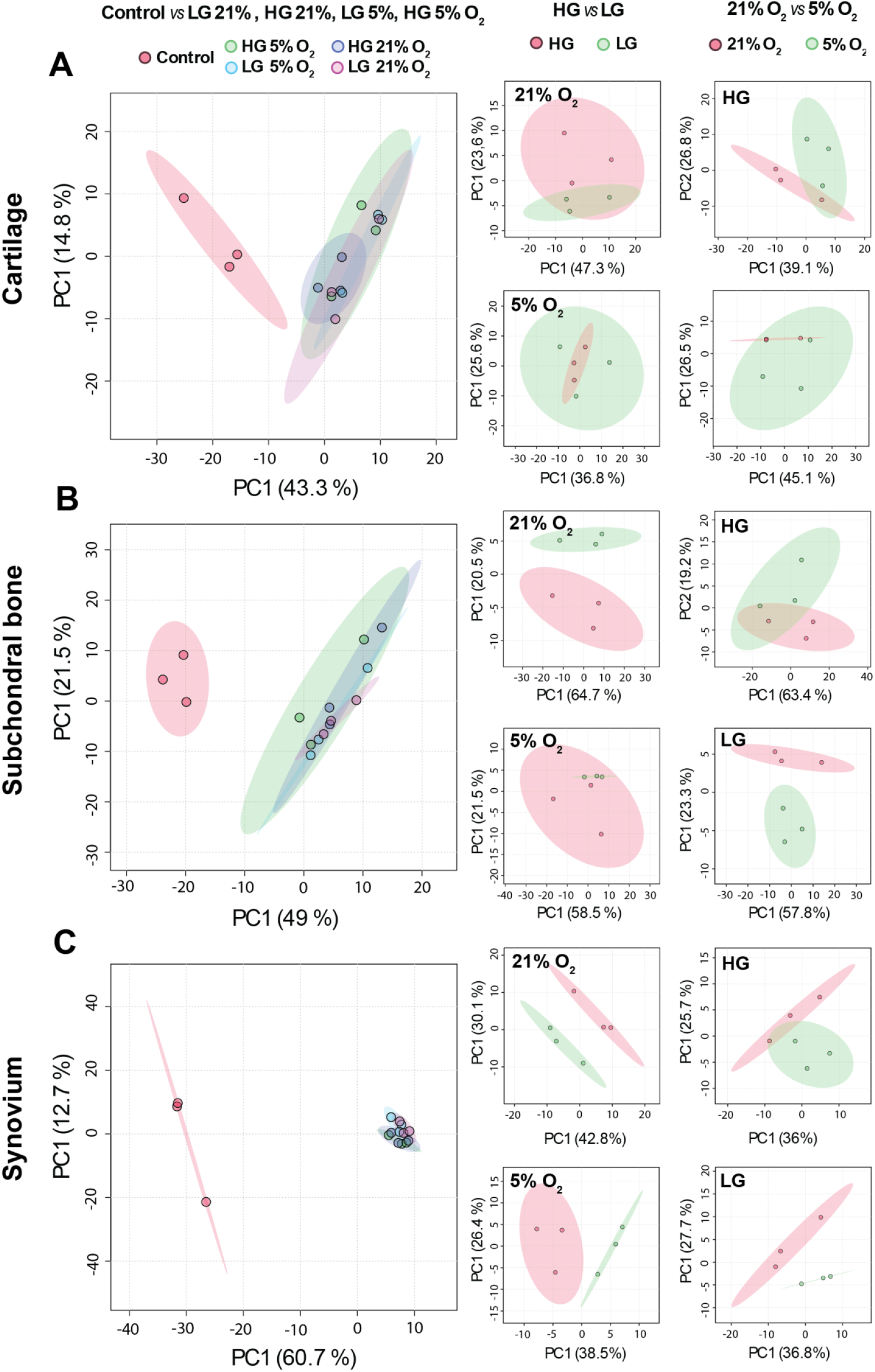
Principle component analysis (PCA) of metabolomics data of (A) cartilage, (B) bone and (C) synovium cultured for 7 days in LG or HG medium under 21% or 5% O_2_ concentration. Fresh tissue is considered as control for each tissue type.

Evaluation of differentially expressed metabolites (log2 fold change [log2FC] > 1) revealed only a limited number of metabolites, with 5–7 in cartilage, 8–14 in bone, and 7–11 in synovium showing significant changes across conditions (supplementary material, Table S1-S3). Among the altered metabolites in cartilage, comparison of HG and LG conditions under 21% O_2_ and 5% O_2_ showed elevated levels of glucose, glucosamine, gluconic acid, while uracil and cAMP were decreased in HG compared to LG (Table S1). Under HG and LG conditions, hypoxanthine and ethyl sulfate were increased whereas 4-pyridoxic acid was decreased in 5% O_2_ compared with 21% O_2_ (Table S1). Among the altered metabolites in bone and synovium, comparison of HG and LG conditions under 21% O_2_ and 5% O_2_ showed increased glucose and gluconic acid levels, and decreased uracil in HG similar to what was observed in cartilage (Table S2, S3). Under both HG and LG conditions, 5% O_2_ downregulated 4-pyridoxic acid, adenine, adenosine, and guanosine, compared with 21% O_2_, whereas ethyl sulfate was upregulated (Table S2, S3).

Pathway analysis of all altered metabolites identified several pathways, mostly represented by a single metabolite, across the experimental comparisons, including 21% and 5% O_2_, as well as the three tissue types (synovium, subchondral bone, and cartilage). However, none of the pathways reached statistical significance after FDR correction. Across all three tissues, glucose conditions modulated pathways related to neomycin, kanamycin, and gentamicin biosynthesis, starch and sucrose metabolism, and galactose metabolism under both 21% and 5% O_2_ (Figure 8A-C). The enrichment of the ‘neomycin, kanamycin, and gentamicin biosynthesis’ pathway was driven by glucose-related metabolites that overlap with KEGG amino-sugar annotations, indicating altered carbohydrate metabolism rather than true antibiotic biosynthesis. Some pathways were specifically regulated by glucose under 21% O_2_, including pantothenate and CoA biosynthesis, the pentose phosphate pathway, β-alanine metabolism, and pyrimidine metabolism, whereas others were modulated only under 5% O_2_, such as arginine and proline metabolism and the pentose phosphate pathway. Across both glucose conditions, pathways associated with purine metabolism, vitamin B6 metabolism, and the biosynthesis of multiple amino acids (valine, leucine, isoleucine, arginine, pyrimidine, alanine, tryptophan, cysteine, aspartate, and glutamate) were commonly regulated by oxygen in all three tissues (Figure 8A-C).

**Figure 8.**
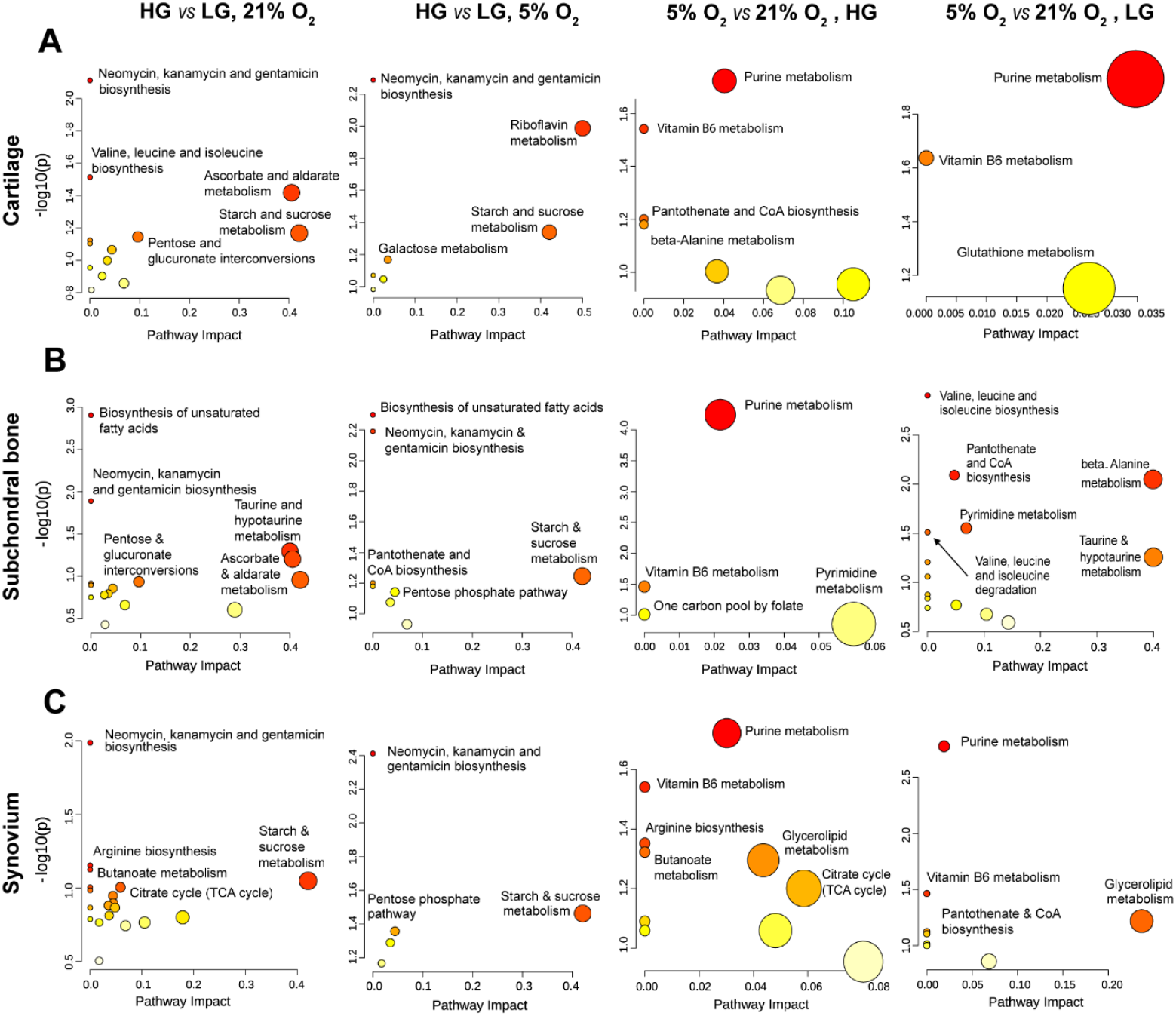
Pathway analysis of the differentially expressed metabolites in (A) cartilage, (B) subchondral bone, and (C) synovium. Pathway analyses are performed separately on the list of metabolites that were differentially expressed in tissues cultured in HG vs LG under either 5% or 21% O_2_. Similarly, the analysis is performed for the differentially expressed metabolites in tissues cultured under 5% vs 21% either in HG or LG condition.

Assessment of glucose levels in cartilage, bone, and synovium, both in fresh control tissues and after 7 days of the culture revealed distinct differences among groups. In cartilage and bone, control samples displayed a broader distribution of normalized log2-transformed glucose levels. Across both tissues, clear differences were observed between HG and LG conditions, with LG groups (at both 21% and 5% O_2_) consistently exhibiting lower glucose levels although it was not significant (p > 0.5). In synovium, glucose levels in the control group were comparable to those observed in samples cultured with LG medium under 5% O_2_ (Figure S2).

## 3 DISCUSSION

One of the most important factors in a successful tissue culture is to maintain cell viability. Investigating cell viability in cartilage, bone and synovium revealed an increase of cell death in cartilage and subchondral bone under LG culture condition independent of oxygen level. This was not observed in synovium and may be explained by its low thickness which allows a more uniform nutrient diffusion across the tissue. However, in cartilage and bone, the high number of dead cells in the central parts of the tissue which are less accessible to nutrients suggests that LG medium fails to support cell maintenance and hence their viability in the entire explant. These results are consistent with a report showing that high glucose level increases cell viability in cartilage [25]. The failure of LG medium to maintain cell viability in deeper regions of cartilage and bone suggests that physiological glucose concentrations alone are insufficient to meet the high metabolic demands of these tissues *ex vivo*. A few factors are likely to contribute to this observation. First, the lack of blood supply in *ex vivo* subchondral bone cultures restricts nutrient transport, leaving cells reliant solely on diffusion from the surrounding medium. Second, the absence of physiological loading further compromises cartilage nutrition, as load-induced fluid advection is omitted in the absence of dynamic loading. Third, static culture conditions fail to replicate *in vivo* dynamics, where synovial fluid is replenished approximately every two hours [26]. Although we replaced the medium daily, this cannot fully mimic *in* vivo nutrient supply. Consequently, HG medium may better sustain cellular metabolic needs and viability over the culture period by providing a larger nutrient reservoir, despite exceeding physiological glucose levels. In the present study, in cartilage, the expression of key anabolic gene *ACAN* was reduced by LG under 21% O_2_, likely reflecting the critical role of glucose as a precursor for glycoprotein and glycosaminoglycan synthesis necessary for extracellular matrix formation. However, the expression of *ACAN*, as well as *COL2* and *COMP* was induced under physiologically relevant oxygen condition (5% O_2_), indicating that oxygen availability influences the regulation of cartilage anabolic genes. This finding aligns with previous studies showing that low oxygen conditions (5% O_2_) enhance the expression of *SOX9* and *ACAN* in human chondrocytes compared to 21% O_2_ [27] supporting a hypoxia-driven anabolic response in cartilage.

Additionally, HG conditions increased the *expression* of several catabolic, hypertrophic, and inflammatory genes in cartilage, including *ADAMTS4, MMP13, COL10A1*, and *IL6*, specifically under 21% of O_2_. Notably, these glucose-dependent differences were not observed under 5% O_2_, where gene expression levels under LG and HG conditions became comparable. Consistent with this, metabolomics analysis showed that LG and HG groups exhibited more similar metabolite profiles at 5% O_2_ than at 21% O_2_, indicating that hypoxia buffers or overrides glucose-mediated metabolic responses. It has been previously reported that both glucose deprivation and hypoxia can independently exert pro-inflammatory effects and induce IL6 expression [28]. However, hypoxia has also been reported to counteract glucose-deprivation–induced IL6 expression in microglia, macrophages, and monocytes [29]. This antagonistic effect appears to occur through Akt/p38 signaling rather than HIF-1/2α pathway. Together with our findings, this supports the presence of a crosstalk between glucose availability and hypoxia pathways, suggesting that transcriptional and metabolic responses to glucose interact with oxygen-sensitive pathways.

In bone and synovium, the effects of glucose and oxygen on gene expression were gene specific. For instance, *ALPL* expression in bone was highly regulated by oxygen but insensitive to glucose availability. This is consistent with multiple studies demonstrating that low oxygen (2–5%) reduces ALPL mRNA, its enzymatic activity, and the formation of mineralized bone nodules [9, 30-32]. *BGLAP*, which encodes osteocalcin, showed a pattern similar to anabolic genes in cartilage: it was downregulated by LG at 21% O_2_ but unaffected by glucose under 5% O_2_. Osteocalcin is a bone-derived endocrine hormone that regulates systemic glucose homeostasis by stimulating insulin secretion in pancreatic cells and increasing peripheral insulin sensitivity. Hyperglycemia in type 2 Diabetes Mellitus Patients has been shown to decrease osteocalcin levels in the blood [33]. Although this appears inconsistent with our *in vitro* observation, the decreased osteocalcin in patients more likely reflects disturbed bone energy metabolism or impaired osteoblast function rather than a direct suppressive effect of glucose on *BGLAP* transcription. *VEGF* expression in bone but not in synovium, was modulated by oxygen. Under LG conditions, 5% O_2_ significantly increased *VEGF*, consistent with the classical hypoxia-induced HIF-1α–mediated stimulation of angiogenic genes [22, 23]. Moreover, previous studies show that low glucose availability results in intracellular ATP depletion which increases *VEGF* mRNA expression [34]. Thus, our result suggests that hypoxia-induced *VEGF* expression is further modulated by cellular energy status, with glucose availability shaping the magnitude of the response.

In synovium, *IL6* followed a pattern similar to cartilage, increasing under HG only at 21% O_2_. This suggests that, unlike bone, cartilage and synovium share a comparable glucose–oxygen interaction profile for *IL6* regulation. However, catabolic markers *MMP1* and *ADAMTS5* displayed divergent responses. *ADAMTS5* expression was reduced by LG in both oxygen levels, whereas *MMP1* expression was decreased under both LG and HG at 5% O_2_. These different patterns indicate a more complex, tissue-specific regulatory landscape in synovium, likely reflecting its heterogeneous cellular composition, including fibroblasts, macrophages, and endothelial cells, in contrast to the relatively homogeneous chondrocyte population in cartilage. Metabolomics further supported this complexity. Synovium showed the strongest metabolic response to glucose and oxygen, with clear PCA clustering between experimental conditions. This clustering was driven primarily by glucose concentration, while in bone the dominant driver was oxygen. Cartilage displayed the weakest clustering, suggesting a more constrained or buffered metabolic plasticity compared with synovium and bone. Owing to the hypoxic nature of cartilage tissue and its dependence on glycolysis, larger metabolic shifts are expected in bone and synovium under low glucose and oxygen availability.

Our data further indicates that glucose and oxygen levels jointly regulate the expression of specific glucose transporters in a tissue-dependent manner. In cartilage, *GLUT1* and *GLUT3* were responsive to both glucose concentration and oxygen levels; in bone, *GLUT1* and *GLUT4* showed similar regulation; and in synovium, *GLUT3* and *GLUT5* were affected. Overall, LG and 5% O_2_ tended to increase *GLUT1* and *GLUT3*, whereas HG and 21% O_2_ increased GLUT4, and LG combined with 5% O_2_ decreased GLUT5 expression. Notably, in all three tissues, *GLUT1* and *GLUT3* were the most abundantly expressed transporters, whereas *GLUT4* and *GLUT5* showed lower baseline levels. Glucose is the primary energy source for mammalian cells, and its availability can fluctuate over time. Cells adapt to such fluctuations through several mechanisms, one of the most important being the modulation of glucose transporter expression. Enhanced *GLUT1* and *GLUT3* expressions in different cell types under hypoxia and through HIF-1α signaling is well known [35]. Furthermore, hypoxia and glucose deprivation act synergistically to upregulate *GLUT1, GLUT3*, and *VEGF* expression in pancreatic cancer cells [36]. *GLUT1, GLUT3*, and *GLUT4* exhibit high affinity for glucose, while *GLUT5* has a high affinity for fructose. In our study, *GLUT5* expression was low in native tissues compared to the other glucose transporters, but its expression increased after ex vivo culture. This likely reflects adaptations to altered nutrient composition in the culture medium, which can shift cellular metabolism. This is supported by the observed substantial differences in metabolite profiles between native tissues and tissues cultured for 7 days, highlighting the metabolic remodeling induced by *ex vivo* conditions.

Metabolomics analysis revealed oxygen-dependent modulation of vitamin B6 metabolism, characterized by consistently reduced levels of 4-pyridoxic acid across cartilage, bone, and synovium under 5% O_2_ compared with 21% O_2_. 4-Pyridoxic acid is a catabolic product of vitamin B6 turnover. The active form of vitamin B6 is Pyridoxal 5′-phosphate (PLP). It is a coenzyme for many amino acid– metabolizing enzymes and is produced from pyridoxine, which is present in DMEM in the form of pyridoxine hydrochloride, via sequential phosphorylation by pyridoxal kinase and oxidation by pyridoxine 5′-phosphate oxidase (PNPO) [37, 38]. The observed reduction in 4-pyridoxic acid indicates diminished degradation of PLP. PNPO has been identified as an oxygen-sensing factor that suppresses lysosomal activity in macrophages under prolonged hypoxia, independent of the HIF pathway [39]. These findings suggest that prolonged low oxygen conditions inhibit PNPO activity and therefore reduce PLP. Consistent with this finding, our data indicates that low oxygen conditions suppress vitamin B6 degradation, and this may be due to the reduced formation of PLP. Furthermore, hypoxic cells rely heavily on amino acid metabolism [40, 41], as reflected by decreased levels of metabolites associated with various amino acid pathways in subchondral bone in our data. This metabolic shift increases demand for PLP-dependent transamination reactions, reinforcing the potential cellular strategy of reducing vitamin B6 breakdown to maintain sufficient PLP levels.

Across all joint tissues, purine metabolism was modulated by oxygen availability. In cartilage, physioxia-induced accumulation of hypoxanthine is consistent with previous reports [42] demonstrating that low oxygen conditions promote a metabolic shift toward glycolysis and increased reliance on nucleotide salvage [43, 44]. In synovium and bone, reduced levels of purine nucleotides and bases (adenosine, guanosine, and adenine) as well as allantoin, the end product of the purine catabolism in mammals, reflect this metabolic shift under physioxic conditions. Under hypoxia, limited ATP availability constrains energy-intensive *de novo* purine synthesis, leading to the accumulation of salvageable purine intermediates as a strategy to conserve cellular energy [44].

Our findings also revealed a distinct tissue-specific pattern of glucose concentration. Comparison of glucose content in fresh tissues with explants cultured for seven days showed that fresh synovium contained glucose levels similar to those cultured under LG culture conditions. This was expected, since LG concentration is the physiological concentration of glucose in synovial fluid similar to blood. In contrast, fresh cartilage and subchondral bone tissues showed glucose levels more comparable to explants maintained in HG medium, although this pattern displayed donor-to-donor variability. This might indicate that HG medium can help maintain glucose levels in cartilage and bone at a level similar to *in vivo* conditions, however these results should be further confirmed using a larger number of donors to better define the native glucose profile of cartilage and subchondral bone.

## 4 CONCLUSION

Taken together, this study demonstrates that glucose concentration and oxygen availability modulate the homeostasis and metabolism of joint tissues in distinct, tissue-specific ways. High-glucose medium appears most suitable for maintaining cartilage and subchondral bone viability under the current tested conditions, whereas synovium exhibited greater sensitivity to variations in its metabolic environment. The differential responses of synovium compared with cartilage and bone highlight the complexity of the joint microenvironment and underscore the challenge of establishing a single *ex vivo* culture condition that is optimal for all joint tissue types. These findings suggest that incorporating physiologically relevant oxygen and glucose gradients into ex vivo joint culture systems, using advanced tissue engineering approaches, may better recapitulate native joint environments in musculoskeletal research. Nevertheless, culturing tissues under physioxic conditions (5% O_2_) helped buffer cartilage and bone responses to nutrient fluctuations, supporting a more stable and physiologically relevant metabolic state. These conditions therefore provide a more optimal environment for joint tissue cultures to study cartilage regeneration and joint-associated disease, where preserving native tissue physiology and minimizing culture-induced artifacts are essential.

## 5 EXPERIMENTAL SECTION/METHODS

### 5.1 Tissue isolation and culture

Bovine stifle joints were obtained from a local abattoir (Metzgerei Angst AG, Zurich, CH) within 48 h of slaughter. Cylindrical osteochondral explants (8 mm in diameter, 6 mm in height) were collected from the femoral groove as previously described [45] (Figure 1). Synovium tissues were collected from the same joint and 8 mm diameter discs were obtained using a biopsy punch (KAI medical, Seki-shi, Gifu Prefecture). Both osteochondral and synovium tissue biopsies were transferred to phosphate-buffered saline (PBS) supplemented with 10% penicillin/streptomycin (P/S, Gibco) for at least 20 min. Every tissue sample was then rinsed once with PBS supplemented with 1% P/S and transferred to a 24-well plate containing 2 ml of Dulbecco’s modified eagle medium (DMEM-HG, 4.5 g/L-glucose; Gibco) supplemented with 10% fetal bovine serum (FBS, Gibco) and 1% P/S. At this time, for metabolomics analysis, tissues were preserved and designated as control fresh tissue. After overnight incubation at 37 °C, 21% O_2_ and 5% CO_2_, co-cultures of osteochondral plugs with synovium tissues were started using either DMEM HG or DMEM LG (1g/L glucose; Gibco) supplemented with 1% P/S, 1% insulin-transferrin-selenium (ITS) + Premix (Corning, Tewksbury, MA, USA), 1% non-essential amino acids (NEAA, Gibco), 50 μg/mL ascorbic acid (Sigma-Aldrich), and 2.5% HEPES (Gibco). To stabilize the osteochondral tissue at the center of the custom-made tissue holders (made of Polyether ether ketone (PEEK), ∼31 mm diameter and ∼10 mm height) and to prevent cell migration from the subchondral bone, 800 µL of 2% agarose gel was applied. After gel solidification, two synovial tissue punches were placed into the tissue holder. Synovium tissues were secured using a 26G needle (Sterican, 0.45 × 25 mm) anchored to the bottom of the holder. The excess portion of the needle was removed using sterile tin snips. At this point, some tissues were preserved and designated as control day 0 for gene expression analysis. *Ex vivo* co-cultures were conducted either in the normal incubator containing 21% O_2_ or in a Hypoxia/Physioxia Workstation (Baker Ruskinn) containing 5% O_2_ for 7 days.

### 5.2 Cell viability assays

Cell viability was evaluated using live/dead staining by Lactate Dehydrogenase (LDH)/ Ethidium homodimer-1 (EthHD1) staining for the osteochondral tissues and calcein AM/ EthHD1 staining for synovium. For this, on day 7 of culture, osteochondral plugs were cut in 2 halves. One half was transferred to cryo-compound and then snap frozen in liquid nitrogen. After cross-sectioning using cryo- tape (Cryofilm type 3C, SECTION-LAB Co) for non-decalcified tissues, 8 µm cuts of osteochondral tissues were stained using LDH/ EthHD1 according to the protocol [46]. Cell viability was then evaluated using Olympus microscopy. For synovium tissue, fresh samples were stained using calcein AM/ EthHD. In brief, freshly prepared solution of 4 mM calcein AM (Sigma-Aldrich) and 2 mM EthHD1 (Sigma-Aldrich) in PBS was used. Synovium tissues were incubated in this solution for 1 hour at 37°C. Cell viability was evaluated using confocal microscopy (LSM 800; Carl Zeiss AG).

Fiji software (1.54g version) was used to analyze the microscopy images. For cartilage sections, the central part and for subchondral bone sections, the central and edge parts of every cross-section were considered for cell counting process as indicated in Figure 2. For synovium tissue, five similar regions of each image were analyzed. The number of dead cells and live cells was counted manually in all images. Four biological replicates were used for semi-quantification and statistical analysis.

### 5.3 Analysis of gene expression by real-time quantitative reverse transcription PCR (RT-qPCR)

Osteochondral tissues from the control group (day 0) and from the experiments (day 7) were first separated into cartilage and bone. Half of the osteochondral explant was used for RNA isolation. All three tissues, including synovium were cut into small pieces using scalpel and were immediately snap-frozen. For RNA isolation, each sample was pulverized using a house-made metal tool and liquid nitrogen. To keep the sample frozen during the pulverization, all the tools were pre-cooled before. Samples were then transferred to a 2-mL Eppendorf tube containing 1 mL TRI reagent (Molecular Research Center) and further homogenized using a 6 mm diameter metal ball and tissue homogenizer for 3 min at 30 Hz frequency for 2-3 times for tissue homogenization. RNA isolation and cDNA synthesis were performed using the RNeasy Mini Kit (Qiagen) and Superscript Vilo RT kit (Thermo Fisher Scientific), respectively. RT-qPCR was then performed using TaqMan Gene Expression Master Mix with QuantStudio Flex 7 instrument. Used primers and probes are listed in Table 1. Gene expression assays from Applied Biosystems were used for additional genes including: *IL8* (Bt03211906_m1), *ALPL* (Bt03244508_m1), *RPLP0* (Bt03218086_m1), *GLUT1* (Bt03215313_m1), *GLUT2* (Bt03258678_m1), *GLUT3* (Bt03259519_gH), *GLUT4* (Bt03215316_m1), *GLUT5* (Bt03258292_m1), *COL1A1* (Bt03225320_g1), *BGLAP* (Bt03213535_g1), and *VEGFA* (Bt03213282_m1).

**Table 1.**
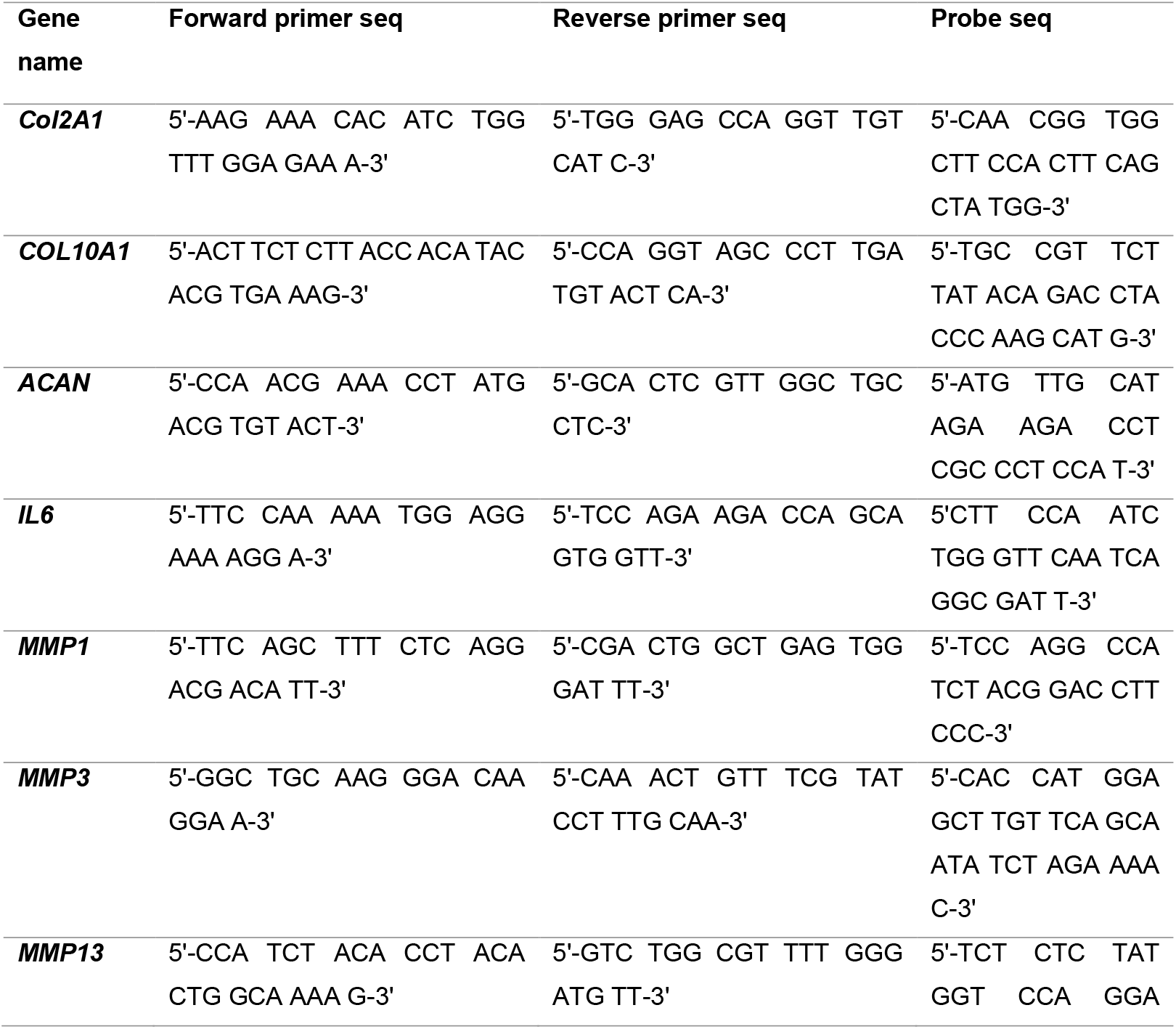
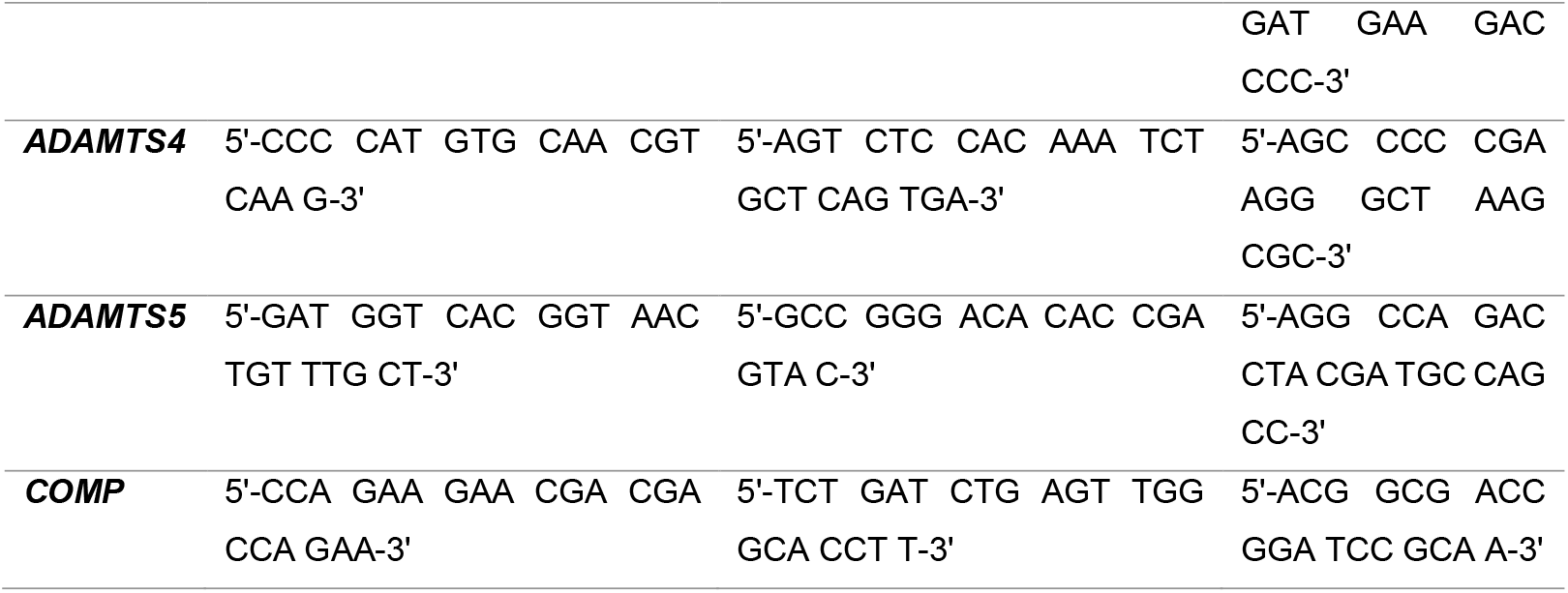
Custom designed primers and probes used for qRT-PCR.

### 5.4 GAG and NO measurement

For evaluation of the glycosaminoglycan (GAG) and nitric oxide (NO) release, conditioned medium was collected every day for 7 days of culture and kept at −20°C till analysis. The amount of GAG released into the medium was determined by the dimethyl-methylene blue (DMMB, Sigma-Aldrich) dye method and analyzed using published methodology protocol [47]. In brief, each sample or chondroitin sulfate standard (Fluka BioChemika) was pipetted into a 96-well plate in duplicate (n = 2), and DMMB solution was added at a 1:4 volume ratio (50 µl and 200 µl respectively). Absorbance was then measured at 535 nm (TECAN® Infinite 200 PRO M Plex, Männedorf, Zürich, CH) and GAG concentration in each sample was determined from a linear calibration curve generated using the chondroitin sulfate standards (chondroitin-4-sulfate sodium salt from bovine trachea, mixture of isomers, Fluka BioChemika). Nitric oxide (NO) released in medium was quantified using the Griess Reagent System (Promega Corporation) according to the manufacturer’s protocol. Absorbance was recorded at 535 nm using the TECAN microplate reader. Nitrite concentrations were calculated using a standard curve generated from sodium nitrite standards.

### 5.5 Histology

For Safranin O/Fast Green staining, cryo-sections, which were prepared as described in the Cell viability assays section, were fixed in 4% formalin (Formafix AG), washed in distilled water (dH2O) to remove the cryo compound, then stained with Weigert’s Haematoxylin solution (Sigma-Aldrich) for 30 min and rinsed in tap water for 10 min. The slides were then stained with 0.02% Fast Green (Fluka #51275) for 6 min and washed in 1% acetic acid for 30 seconds. Then they were stained with 0.1% Safranin-O (Sigma-Aldrich) for 12 min. Subsequently, samples were rinsed with dH_2_O. After dehydration with increasing concentrations of ethanol, samples were transferred to xylene and cover-slipped with Eukitt mounting medium (Sigma-Aldrich).

### 5.6 Sample preparation for LC-MS analysis

Untargeted metabolomics was performed by the Functional Genomics Center Zurich (ETH Zurich/University of Zurich, Switzerland) on *Bos taurus* bones, cartilage and synovium samples, separately. Frozen tissues were cryopulverized and the polar metabolites extracted with ice-cold methanol:water (80:20, v/v). Fresh tissues were used as control. Homogenates were centrifuged for 10 min at 14000 rpm/4°C and metabolite extracts in supernatant were transferred to clean test tubes. Prior to measuring, extracts were dried under nitrogen flow and reconstituted in injection buffer (90% acetonitrile). The solution was vortexed and centrifuged for 10min at 14000 rpm/4°C and 60 mL of the supernatants were transferred to glass vials with narrowed bottom (Total Recovery Vials, Waters) for liquid chromatography and mass spectrometry (LC-MS) injection. In addition, method blanks, quality control (QC) standards, and pooled samples were prepared in the same way to serve as quality control for the measurements.

### 5.7 LC-MS analysis

Polar metabolites were separated on a Thermo Vanquish Horizon Binary Pump equipped with Waters Premier BEH Amide column (150 mm x 2.1 mm), applying a gradient of 10 mM ammonium bicarbonate in 5% acetonitrile (A) and 10 mM ammonium bicarbonate in 95% acetonitrile (B) from 99% B to 30% B over 12 min. The injection volume was 4 mL. The flow rate was 0.4 uL/min with a column temperature of 40°C and autosampler temperature of 5°C.

The LC was coupled to Thermo Exploris 480 mass spectrometer by a HESI source. MS1 (molecular ion) and MS2 (fragment) data were acquired using negative polarization and Full MS / dd-MS^2^ (Top5) over a mass range of 70 to 1050 m/z at MS1 resolution of 60000 and MS2 resolution of 7500.

### 5.8 Untargeted Metabolomics Data analysis

The metabolomics data set was evaluated in an untargeted fashion with Compound Discoverer software (Thermo Scientific). The modular data analysis workflow includes spectra selection, retention times alignment, compound detection and grouping, gap filling, background filtering, and normalization (data are constant median normalized). mzCloud and mzVault were used to score fragmentation patterns and assign MS2-based identities to the features. A filtering process was performed, leading to the manually annotated compound table, where each feature is annotated with the highest level of confidence. Filtering parameters used were the following: Signa/noise > 3, mzCloud or mzVault match > 50, ppm mass error within +/- 5 ppm., match with in-house developed MS1_RT library within +/- 10 sec., chromatographic peak and MS2 spectra quality. Quality controls were run on pooled QC samples and reference compound mixtures to determine technical accuracy and stability.

Prior to downstream data analysis, metabolite intensity values for each tissue sample were normalized to the total protein content (determined on the protein pellet resulting from polar metabolites extraction). The resulting annotated metabolite table was subsequently imported into MetaboAnalyst version 6.0 for downstream analysis. To improve data comparability and reduce heteroscedasticity, metabolite intensities were log-transformed, followed by autoscaling (mean-centering). This preprocessing approach ensured that metabolites with different dynamic ranges contributed equally to multivariate analyses. The principal component analysis (PCA) was performed separately for each tissue type to visualize global metabolic differences and clustering patterns between experimental groups. PCA was further used as an exploratory tool to assess overall variability and identify potential group-level separation in metabolomic profiles. Student’s t-test was used to evaluate differences in metabolite levels between experimental groups. Metabolites with an FDR-adjusted p-value < 0.05 and log2fold change >1 were considered significantly differentially regulated between conditions. Metabolites identified as significantly altered were subsequently subjected to pathway enrichment analysis using the Functional Analysis module in MetaboAnalyst. Statistical significance for pathway enrichment was determined using an FDR-corrected threshold of p < 0.05.

### 5.9 Statistical analysis

Quantitative data are presented as mean ± standard deviation. Statistical analyses were performed using GraphPad Prism 10. Differences between groups were assessed using two-way ANOVA followed by Tukey’s post hoc multiple comparison test, unless otherwise specified in the figure legends. A p-value < 0.05 was considered statistically significant.

## ACKNOWLEDGMENTS

This work was supported by the AO Foundation. Mobility and research exchange activities for Dr. Jovana Zvicer to visit the AO Research Institute were supported by the European Union’s Horizon 2020 research and innovation programme under grant agreement No. 952033 (ExcellMater – Twinning to Excel Materials Engineering for Medical Devices) and by the NetwOArk COST Action (CA21110), funded by COST (European Cooperation in Science and Technology). We would like to thank Dr. Jacek Wychowaniec for his helpful feedback on the manuscript.

## DATA AVAILABILITY STATEMENT

The data that supports the metabolomics findings of this study are available from the corresponding author upon request.

