## Supplementary material for "Toward Standardized Ex Vivo Joint Models: Impact of Glucose and Oxygen Levels for Enhanced Tissue Maintenance": Table S1

**Table S1:** Differentially regulated metabolites in cartilage co-cultured with bone and synovium under 21% and 5% $O_2$ and high glucose (HG) and low glucose (LG) medium for 7 days.

| **HG vs LG, 21% $O_2$** | | | | |
|---|---|---|---|---|
| **Metabolites** | **FC** | **log2(FC)** | **raw.pval** | **-LOG10(p)** |
| Uracil | 0.24091 | -2.0534 | 0.024236 | 1.6155 |
| THREONINE | 0.25692 | -1.9606 | 0.060481 | 1.2184 |
| UDP-N-acetylglucosamine | 2.0025 | 1.0018 | 0.087007 | 1.0604 |
| N-ACETYLSERINE | 2.4905 | 1.3164 | 0.032678 | 1.4858 |
| Glucuronic acid | 4.4036 | 2.1387 | 0.039211 | 1.4066 |
| Gluconic acid | 8.4046 | 3.0712 | 0.000438 | 3.3587 |
| Glucose | 9.2894 | 3.2156 | 0.00414 | 2.383 |
| **HG vs LG, 5% $O_2$** | | | | |
| cAMP | 0.23195 | -2.1081 | 0.014191 | 1.848 |
| MALEATE | 0.38528 | -1.376 | 0.05722 | 1.2425 |
| Riboflavin | 0.48773 | -1.0359 | 0.054337 | 1.2649 |
| GLUCOSAMINE | 2.5663 | 1.3597 | 0.006135 | 2.2122 |
| Creatine | 2.9909 | 1.5806 | 0.035561 | 1.449 |
| Glucose | 6.1875 | 2.6294 | 0.013849 | 1.8586 |
| **HG, 5% $O_2$ vs 21% $O_2$** | | | | |
| 4-Pyridoxic acid | 0.05244 | -4.2532 | 0.001978 | 2.7038 |
| Uric acid | 0.23352 | -2.0984 | 0.048272 | 1.3163 |
| PE 36:3 | 0.2833 | -1.8196 | 0.084566 | 1.0728 |
| 4-Hydroxybenzaldehyde | 3.9644 | 1.9871 | 0.069715 | 1.1567 |
| Uracil | 4.2057 | 2.0723 | 0.072206 | 1.1414 |
| ETHYL SULFATE | 6.2389 | 2.6413 | 0.003194 | 2.4956 |
| Hypoxanthine | 16.807 | 4.071 | 0.006851 | 2.1642 |
| **LG, 5% $O_2$ vs 21% $O_2$** | | | | |
| 4-Pyridoxic acid | 0.091929 | -3.4433 | 0.021668 | 1.6642 |
| Cytidine 2',3'-cyclic monophosphoric acid | 0.38716 | -1.369 | 0.050428 | 1.2973 |
| Guanine | 5.7683 | 2.5282 | 0.009296 | 2.0317 |
| Hypoxanthine | 7.7285 | 2.9502 | 0.078904 | 1.1029 |
| ETHYL SULFATE | 7.977 | 2.9959 | 0.008906 | 2.0503 |
