## Supplementary material for "Toward Standardized Ex Vivo Joint Models: Impact of Glucose and Oxygen Levels for Enhanced Tissue Maintenance": Table S2

**Table S2:** Differentially regulated metabolites in subchondral bone co-cultured with cartilage and synovium under 21% and 5% $O_2$ and high glucose (HG) and low glucose (LG) medium for 7 days.

| HG vs LG, 21% $O_2$ | | | | |
|---|---|---|---|---|
| **Metabolites** | **FC** | **log2(FC)** | **raw.pval** | **-LOG10(p)** |
| N-Oleoyl-Phenylalanine | 0.21973 | -2.1862 | 0.007756 | 2.1103 |
| Docosahexaenoic acid | 0.28619 | -1.8049 | 0.097043 | 1.013 |
| Uracil | 0.30717 | -1.7029 | 0.060919 | 1.2152 |
| Eicosapentaenoic acid | 0.32653 | -1.6147 | 0.074021 | 1.1306 |
| Arachidonic acid | 0.36249 | -1.464 | 0.029427 | 1.5313 |
| Oleoyl-L-lysophosphatidic acid | 0.39282 | -1.3481 | 0.077693 | 1.1096 |
| Xanthine | 0.39468 | -1.3412 | 0.027004 | 1.5686 |
| Cytidine 2',3'-cyclic monophosphoric acid | 0.43005 | -1.2174 | 0.075814 | 1.1203 |
| Hypotaurine | 0.47437 | -1.0759 | 0.083897 | 1.0763 |
| O-Acetylcarnitine | 2.8516 | 1.5118 | 0.005345 | 2.272 |
| Glucuronic acid | 2.8886 | 1.5304 | 0.055454 | 1.2561 |
| Mannitol | 5.8621 | 2.5514 | 0.011107 | 1.9544 |
| Gluconic acid | 6.448 | 2.6889 | 0.023518 | 1.6286 |
| Glucose | 8.7496 | 3.1292 | 0.00373 | 2.4283 |
| **HG vs LG, 5% O2** | | | | |
| 6-Ketoprostaglandin F1 | 0.3622 | -1.4651 | 0.003925 | 2.4062 |
| Eicosapentaenoic acid | 0.37551 | -1.4131 | 0.017983 | 1.7451 |
| Uracil | 0.38288 | -1.385 | 0.061884 | 1.2084 |
| N-Oleoyl-Phenylalanine | 0.39652 | -1.3345 | 0.005011 | 2.3001 |
| OLEATE | 0.41952 | -1.2532 | 0.072024 | 1.1425 |
| Oleoyl-L-lysophosphatidic acid | 0.48093 | -1.0561 | 0.050449 | 1.2972 |
| Gluconic acid | 3.94 | 1.9782 | 0.024241 | 1.6154 |
| Glucose | 6.5626 | 2.7143 | 0.005715 | 2.243 |
| **HG, 5% $O_2$ vs 21% $O_2$** | | | | |
| 4-Pyridoxic acid | 0.02733 | -5.1934 | 0.001359 | 2.8669 |
| Adenine | 0.081189 | -3.6226 | 0.016001 | 1.7958 |
| Adenosine | 0.15999 | -2.644 | 0.031398 | 1.5031 |
| Inosine | 0.21202 | -2.2377 | 0.016448 | 1.7839 |
| Uridine | 0.23317 | -2.1005 | 0.058965 | 1.2294 |
| cAMP | 0.31547 | -1.6644 | 0.088069 | 1.0552 |
| Mannitol | 0.32654 | -1.6147 | 0.096037 | 1.0176 |
| Guanosine | 0.32928 | -1.6026 | 0.01599 | 1.7962 |
| O-acetylcarnitine | 0.42825 | -1.2235 | 0.023368 | 1.6314 |
| N-Oleoyl-Phenylalanine | 2.2618 | 1.1775 | 0.00533 | 2.2733 |
| Ethyl sulfate | 3.9566 | 1.9842 | 0.026346 | 1.5793 |
| **LG, 5% $O_2$ vs 21% $O_2$** | | | | |
| 4-Pyridoxic acid | 0.081231 | -3.6218 | 0.008296 | 2.0811 |
| Uracil | 0.2581 | -1.954 | 0.022509 | 1.6476 |
| Allantoin | 0.37127 | -1.4294 | 0.073715 | 1.1324 |
| Guanosine | 0.38531 | -1.3759 | 0.005856 | 2.2324 |

| | | | | |
|---|---|---|---|---|
| L-Alanine | 0.41558 | -1.2668 | 0.057742 | 1.2385 |
| Methionine | 0.41793 | -1.2587 | 0.046715 | 1.3305 |
| Beta-alanine | 0.4301 | -1.2173 | 0.08368 | 1.0774 |
| Isoleucine | 0.4609 | -1.1175 | 0.093091 | 1.0311 |
| Leucine | 0.47486 | -1.0744 | 0.061062 | 1.2142 |
| Pseudouridine | 0.47951 | -1.0604 | 0.002421 | 2.6159 |
| Hypotaurine | 0.49365 | -1.0185 | 0.094056 | 1.0266 |
| Tryptophan | 0.49909 | -1.0026 | 0.030129 | 1.521 |
| Ethyl sulfate | 7.9572 | 2.9923 | 0.001433 | 2.8439 |
