## Supplementary material for "Toward Standardized Ex Vivo Joint Models: Impact of Glucose and Oxygen Levels for Enhanced Tissue Maintenance": Table S3

**Table S3:** Differentially regulated metabolites in synovium co-cultured with cartilage and bone under 21% and 5% $O_2$ and high glucose (HG) and low glucose (LG) medium for 7 days.

| **HG vs LG, 21% $O_2$** | | | | |
|---|---|---|---|---|
| **Metabolites** | **FC** | **log2(FC)** | **raw.pval** | **-LOG10(p)** |
| 4-Hydroxybenzaldehyde | 0.064638 | -3.9515 | 0.04573 | 1.3398 |
| Uracil | 0.20494 | -2.2867 | 0.004539 | 2.343 |
| Adenine | 0.38519 | -1.3764 | 0.092891 | 1.032 |
| Indoxyl sulfate | 0.39087 | -1.3552 | 0.066435 | 1.1776 |
| PE 36:3 | 0.39326 | -1.3464 | 0.034306 | 1.4646 |
| Cystathionine | 0.46416 | -1.1073 | 0.025409 | 1.595 |
| Proline | 0.47847 | -1.0635 | 0.096215 | 1.0168 |
| Oxoglutaric acid | 6.4519 | 2.6897 | 0.000178 | 3.7492 |
| Gluconic acid | 16.248 | 4.0222 | 0.007504 | 2.1247 |
| Mannitol | 21.082 | 4.3979 | 0.010567 | 1.976 |
| Glucose | 27.886 | 4.8015 | 0.000565 | 3.2479 |
| **HG vs LG, 5% $O_2$** | | | | |
| Methyl beta-D-galactoside | 0.27902 | -1.8416 | 0.063715 | 1.1958 |
| N-Acetyl-L-alanine | 2.2305 | 1.1574 | 0.015312 | 1.815 |
| Methionine sulfoxide | 2.5382 | 1.3438 | 0.087481 | 1.0581 |
| D-Alanine | 3.1097 | 1.6368 | 0.06285 | 1.2017 |
| Mannitol | 14.507 | 3.8587 | 0.032246 | 1.4915 |
| Gluconic acid | 20.188 | 4.3354 | 0.006813 | 2.1667 |
| Glucose | 20.233 | 4.3386 | 0.00021 | 3.6774 |
| **HG, 5% $O_2$ vs 21% $O_2$** | | | | |
| 4-Pyridoxic acid | 0.13107 | -2.9316 | 0.051091 | 1.2917 |
| Adenosine | 0.27024 | -1.8877 | 0.066161 | 1.1794 |
| Oxoglutaric acid | 0.29825 | -1.7454 | 8.14E-05 | 4.0893 |
| 5,5-dimethyl-2-{[(2-phenylacetyl)amino]methyl}-1,3-thiazolane-4-carboxylic acid | 2.2363 | 1.1611 | 0.001596 | 2.797 |
| Xanthine | 2.3017 | 1.2027 | 0.08963 | 1.0475 |
| 4-Oxoproline | 2.3391 | 1.2259 | 0.005165 | 2.2869 |
| Quinolinecarboxylic acid | 2.8288 | 1.5002 | 0.034318 | 1.4645 |
| D-Alanine | 3.8629 | 1.9497 | 0.028525 | 1.5448 |
| Glycerol 3-phosphate | 4.5189 | 2.176 | 0.06977 | 1.1563 |
| Ethyl sulfate | 10.106 | 3.3371 | 0.008218 | 2.0853 |
| **LG, 5% $O_2$ vs 21% $O_2$** | | | | |
| 4-Pyridoxic acid | 0.063457 | -3.9781 | 0.013486 | 1.8701 |
| Adenosine | 0.23005 | -2.12 | 0.030318 | 1.5183 |
| Guanosine | 0.23009 | -2.1198 | 0.063591 | 1.1966 |
| Adenine | 0.24346 | -2.0382 | 0.028188 | 1.5499 |
| 6-Ketoprostaglandin F1 | 0.24847 | -2.0089 | 0.085871 | 1.0662 |
| Methyl beta-D-galactoside | 0.31279 | -1.6767 | 0.071452 | 1.146 |
| Uracil | 0.3457 | -1.5324 | 0.060529 | 1.218 |
| Glycerol | 5.8899 | 2.5582 | 0.085104 | 1.0701 |
| Ethyl sulfate | 9.9798 | 3.319 | 0.000763 | 3.1173 |
