## Supplementary material for "Toward Standardized Ex Vivo Joint Models: Impact of Glucose and Oxygen Levels for Enhanced Tissue Maintenance": Figure S1

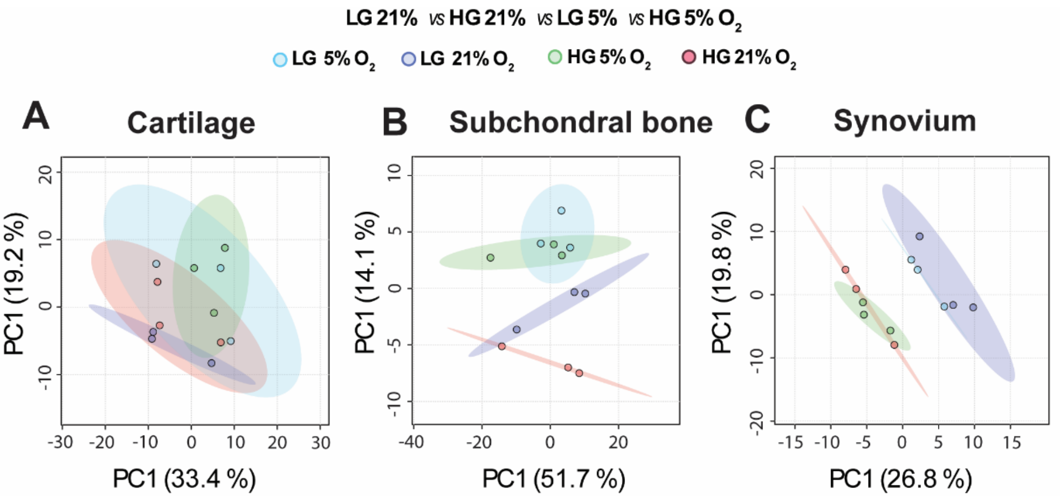


**Figure S1:** Principle component analysis (PCA) of metabolomics data of (A) cartilage, (B) bone and (C) synovium cultured for 7 days in LG or HG medium under 21% or 5% O2 concentration.
