## Supplementary material for "Toward Standardized Ex Vivo Joint Models: Impact of Glucose and Oxygen Levels for Enhanced Tissue Maintenance": Figure S2

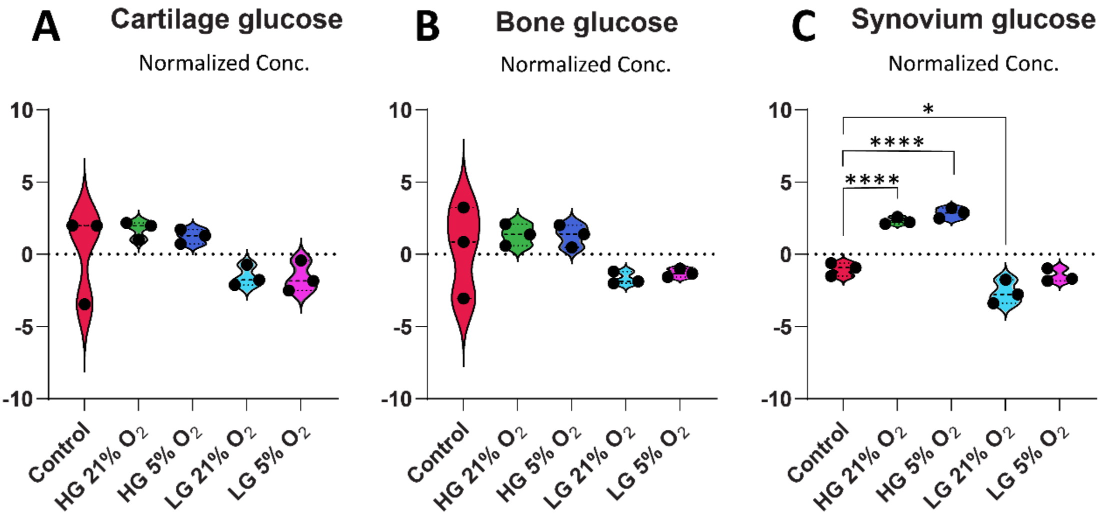


**Figure S2**: Normalized log2-transformed glucose levels in fresh and 7-day cultured tissues in (A) cartilage, (B) subchondral bone, and (C) synovium. Each dot represents an individual biological replicate. N=3. Control is the fresh tissue. Statistical analysis was performed using one-way ANOVA, with significance levels indicated as *$p < 0.05$, ****$p < 0.0001$.
